# Spatial molecular imaging of the human type 2 diabetic islet

**DOI:** 10.1101/2023.01.04.519955

**Authors:** Grant R. Kolar, David Ross, Emily E. Killingbeck, Willis K. Samson, Gina L. C. Yosten

## Abstract

The islets of Langerhans are complex micro-organs comprised of multiple cell types essential for the maintenance of glucose homeostasis. The endocrine cell types of the islets engage in intimate, intercellular communication that is necessary for normal secretory activity. Disruption of this intercellular communication, which is at least partially dependent on the spatial organization of individual islets, leads to secretory dysfunction and exacerbation of the symptomology of disease states such as type 2 diabetes. However, the molecular determinants of the mechanisms underlying disrupted intercellular communication remain incompletely understood. Herein we describe the utilization of CosMx™ Spatial Molecular Imaging (SMI) to interrogate transcriptomic changes associated with the transition from the obese, prediabetic state to overt type 2 diabetes. Using SMI, we verified previously reported findings regarding islet composition in the obese and type 2 diabetic states, including loss of beta cells and expansion of alpha cell mass. In addition, we identified changes in the islet neighborhood that have implications for the function of islet endocrine cells. In particular, we identified a subset of alpha cells oriented in the periphery of the islets that appear to exhibit a transcriptomic profile suggestive of de-differentiation toward a beta cell-like transcriptome-type. To our knowledge, this is the first study utilizing spatial molecular imaging to investigate single-cell transcriptomes of individual islets. Further exploration of the intersection of islet architecture and gene expression using spatial technologies is expected to yield novel insights into the mechanisms underlying the development and progression of metabolic diseases like type 2 diabetes.

## MAIN TEXT

Intercellular communication is essential for the normal functioning of multicellular organ systems. This is particularly true of the islets of Langerhans, which are dependent on complex paracrine and autocrine interactions for the maintenance of hormonal secretory activity^1^. The islets of Langerhans are micro-organs comprised of heterogeneous cell populations but are dominated by the presence of alpha, beta, and delta cells^1^. Removal or dysfunction of any of these major islet cell types results in impairments in the biological activity of the remaining cells^1^. For example, loss of beta cell secretory activity as occurs in the setting of type 2 diabetes (T2D)^2^ leads to aberrations in counterregulation mediated by the alpha cell^1^ and paradoxical release of glucagon even under high glucose conditions (e.g., postprandially)^3^ mediated in part by impairments in somatostatin release from the delta cells^4,5^. Although studies utilizing single-cell RNA sequencing (scRNA-Seq) analysis of islet cell subtypes have provided important insights into the deleterious biological changes associated with T2D^6^, the utility of these methods is limited by their lack of spatial and anatomical context^7–10^. To overcome this limitation, we used CosMx Spatial Molecular Imaging (SMI) (NanoString^®^ Technologies, Seattle, WA)^11^ to evaluate anatomically-relevant transcriptomic changes in islets from patients with T2D and matched non-diabetic patients.

We collected pancreas tissues from three individuals with clinically-confirmed T2D, three individuals with marked obesity but without clinical T2D (metabolically-healthy obese, MHO), and three individuals with no history of obesity or T2D (Normal). Patient characteristics are detailed in **Supplementary Table 1**. Tissues from the tail of the pancreas were rapidly preserved in formalin, embedded in paraffin, and processed for SMI. For each patient sample, an average of 20 fields of view (FOV) were selected based on the relative abundance of histologically-identified islets (**Fig. 1A**); each FOV represented an average of 6,000 islet cells that were subsequently evaluated using an array of probes representing 990 human genes (**Supplemental Table 2**). FOVs corresponding to the H&E image of the pancreas were readily identifiable via the CosMx viewer (**Fig. 1B**). Initial unsupervised clustering of cells based on gene expression using the Leiden algorithm^12^ revealed separate endocrine, exocrine, and immune populations. (**Fig. 1C**). Supervised cell typing using the Insitutype^28^ method and guided by reference profiles from scRNA-seq experiments^13^ achieved more granular cell type classifications that matched immunohistochemically-identified cell types—identified with post hoc immunofluorescent imaging of the tissues—and resolved to real space. (**Fig. 1D** and **Supplemental Videos**). Importantly, UMAP analysis^14,15^ computed using immunohistochemistry-based cell identification (**Fig. 1E, right panel**) matched that of cell identification via Insitutype (**Fig. 1E, left panel**). Thus, this methodology allows for cell type identification with high fidelity. Additional quality control and run metrics are detailed in **Supplemental Table 3**.

**Figure 1:**
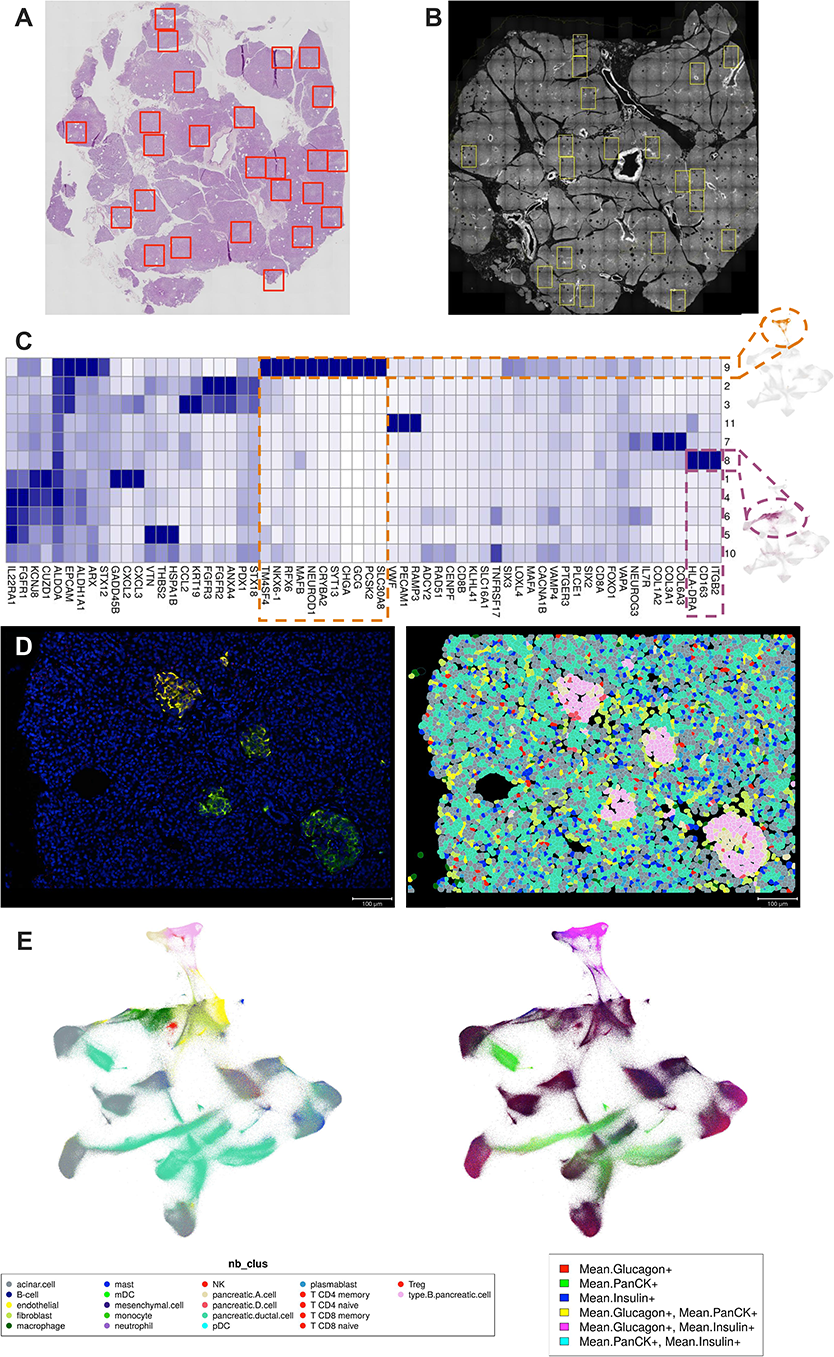
CosMx SMI allows transcript and protein expression correlation separating the exocrine and endocrine pancreas. H&E image of pancreas section showing an overlay of selection ROIs (red) (A) corresponding to a auto-fluorescent image overview of an adjacent serial tissue section on the CosMx platform with the same ROIs selected (yellow) (B). Expression of genes typical of endocrine and exocrine cells are efficiently segregated by clustering to different regions of a UMAP representation (C). Expression of RNA transcripts associated with cell identity allows identification of cell types, segmented with a customized Cellpose algorithm, in a selected ROI (D, right) particularly highlighting alpha cells (tan) and beta cells (pink) that correspond to these same types as identified by immunofluorescence with glucagon (red) and insulin (green) (D, left). UMAP space projection (E) for cell clustering based on transcript expression (left) and overlaid with immunofluorescence expression in the same cells (right) shows clear segregation of endocrine from the exocrine pancreas.

Analysis of islets from patients with T2D by immunohistochemistry^16^, multi-spectral imaging mass spectrometry^17^, and multi-omics techniques^18^ have consistently demonstrated disruptions in islet cell composition and morphology compared to islets from non-diabetic individuals. We therefore evaluated potential differences in islet composition between patient groups following confirmation of accurate cell identification using SMI. Consistent with previous reports^1,16^, islets from non-obese, non-diabetic control patients were composed primarily of endocrine cells (i.e. alpha, beta, and delta cells), with an average of 89.5% of islet cells identified as one of the three major endocrine cell types (**Fig. 2A,B**). Islets from both MHO and T2D individuals exhibited reductions in endocrine cells (75.8% and 76.2% endocrine cells, respectively). Beta cell mass, measured as a percentage of total islet cells, was also reduced in MHO (49.5%) and T2D (51.1%) islets compared to islets from non-diabetic, non-obese individuals (65.4%). There have been mixed reports regarding changes to alpha cell mass in obesity and T2D, with some reports indicating alpha cell hypertrophy in the setting of obesity or T2D^16,19,20^ and other groups reporting no change in alpha cell mass^21^. We noted a slight increase in the percentage of alpha cells in the islets of MHO (23.4%) and T2D (21.3%) individuals compared to lean control patients (19.7%). The beta:alpha cell ratio in islets from MHO patients was significantly higher than in islets from normal and T2D patients (**Fig. 2C**). However, no significant differences in the beta:delta cell ratio (**Fig. 2D**) or the alpha:delta cell ratio (**Fig. 2E**) were observed between groups. Thus, spatial molecular imaging is able to corroborate previous findings regarding changes in islet composition in the setting of obesity or T2D.

**Figure 2:**
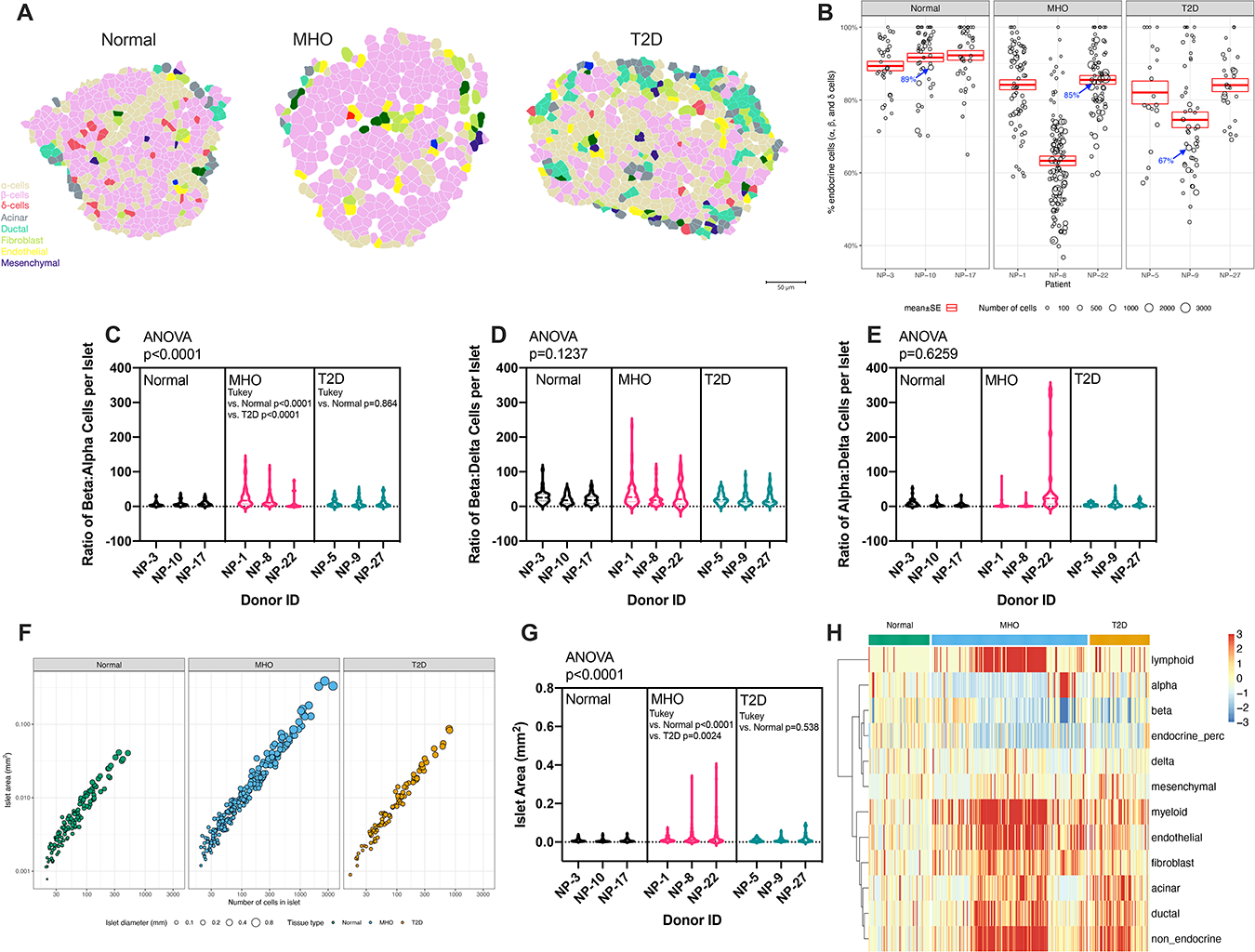
Islet morphology and cell composition differ between experimental groups. A) Examples of single islets in non-obese, non-diabetic (normal) patients, MHO (metabolically healthy obese) patients, and obese donors with type 2 diabetes (T2D). Islets are shown with cell delineation and phenotype identified by corresponding colors. B) Bubble plot showing percentages of endocrine cells (alpha, beta, and delta cells) for individual islets for each donor. Each bubble represents a single islet. Larger bubbles indicate larger islets, in terms of cell counts per islet (see legend underneath graph). Blue arrows with the associated percentage correspond to the islets depicted in (A). C) Violin plot of the beta:alpha cell ratios of individual islets for each donor. The beta:alpha cell ratio was significantly increased in islets from MHO donors compared to normal and T2D donors. Violin plots are shown depicting beta:delta (D) and alpha:delta (E) cell ratios for individual islets from each donor. No significant differences in these measures were observed. F) Bubble plot comparing the number of cells per islet (x axis) to the islet area (y axis) for individual islets for each experimental group. Each bubble represents a single islet. Bubble size indicates islet diameter, such that larger bubbles represent islets with larger diameters (see legend below figure). G) Violin plot representing the area of individual islets for each donor for the three experimental groups. Islets from MHO patients were significantly larger than islets from normal donors or donors with T2D. H) Heatmap showing changes in islet composition compared to islets from normal donors. Columns represent single islets. Blue represents a decrease in the number of a cell type, red indicates an increase in the number of a particular cell type, and yellow indicates no change in composition. Data in C, D, E, and G were analyzed by one way ANOVA with Tukey post-hoc analysis for multiple comparisons. Exact p values are indicated in the figures.

Obesity is an independent risk factor for the development of T2D^22,23^; thus, longstanding obesity is often associated with prediabetes and could be regarded as a transition period from normal metabolic function to T2D. In the obese, prediabetic state, insulin resistance predominates and possibly leads to islet hypertrophy^24,25^. Comparison of the area of individual islets with the number of cells for each islet revealed similar slopes between islets from normal, MHO, and T2D patients (**Fig. 2F**). However, the overall area of the islets appeared to be larger in MHO patients compared to islets from the other groups (**Fig. 2F**). We then specifically assessed the area of islets for each group and found that islets from MHO individuals were 3.12 and 1.95 times larger than that of normal and T2D islets, respectively (**Fig. 2G**). Although islets from T2D patients exhibited an area 1.6 times that of normal islets on average, this difference did not attain statistical significance. Given that islets from MHO and T2D patients exhibited reduced percentages of endocrine cell types, yet larger overall area compared to islets collected from normal, non-obese, non-diabetic individuals, we performed global composition analysis to investigate which cell types within the islets were undergoing the most significant changes in terms of the abundance within the islets. Consistent with previous reports^16^, our data indicated that islets from MHO and T2D patients contained reduced endocrine cell numbers, particularly beta cells (**Fig. 2H**). The percentages of non-endocrine cell types in MHO and T2D islets appeared to be generally expanded compared to normal islets, particularly in islets from MHO patients. For example, lymphoid cells were extensively expanded in islets from MHO patients compared to islets from non-diabetic, non-obese patients (**Fig. 2H, first row**); however, the percentage of lymphoid cells in islets from T2D donors resembled that of non-diabetic, non-obese individuals. These data suggest that in the obese state, overall islet cell-type expansion occurs, perhaps in an attempt to compensate for insulin resistance^22,23^, and supports the hypothesis that the MHO condition represents a transitional state from metabolic instability to T2D.

Differences in the organization of the islet neighborhood in the setting of T2D have been noted previously by Wu and colleagues^17^. In particular, Wu et al. determined that CD8+ lymphocytes are more closely associated with beta cells in islets isolated from donors with T2D compared to islets from non-diabetic patients, while in the non-diabetic state macrophage and beta cell interactions are more likely^17^. We therefore hypothesized that when we investigated the spatial association of islet cell types stratified by patient type, lymphocytes would be spatially positioned close to beta cells in the MHO and T2D states but not in the islets from normal donors. Surprisingly, our dataset among the three major endocrine cell types demonstrated that lymphoid cells were most closely associated with delta cells uniquely in islets from MHO patients (**Fig. 3**). In addition, we did not observe any differences in the proximity of myeloid cells to beta cells across the three patient groups. The discrepancies between our findings and that of Wu et al. could be due to multiple experimental factors. For instance, our dataset consisted of only 3 individual patients for each group, and thus an expansion of our dataset could yield different results. Whereas Wu et al. utilized larger group sizes, their datasets did not include a non-obese, non-diabetic patient group^17^. Also, cell identification and segmentation in their study depended on the antibody-based identification of 34 proteins, while in our study, cells were identified by mRNA expression of 990 genes (**Supplemental Table 2**). Expansion and further development of both multi-spectral imaging and spatial molecular imaging to interrogate islet biology will likely reveal fundamental information regarding the molecular biology underlying the progression of prediabetes to T2D.

**Figure 3:**
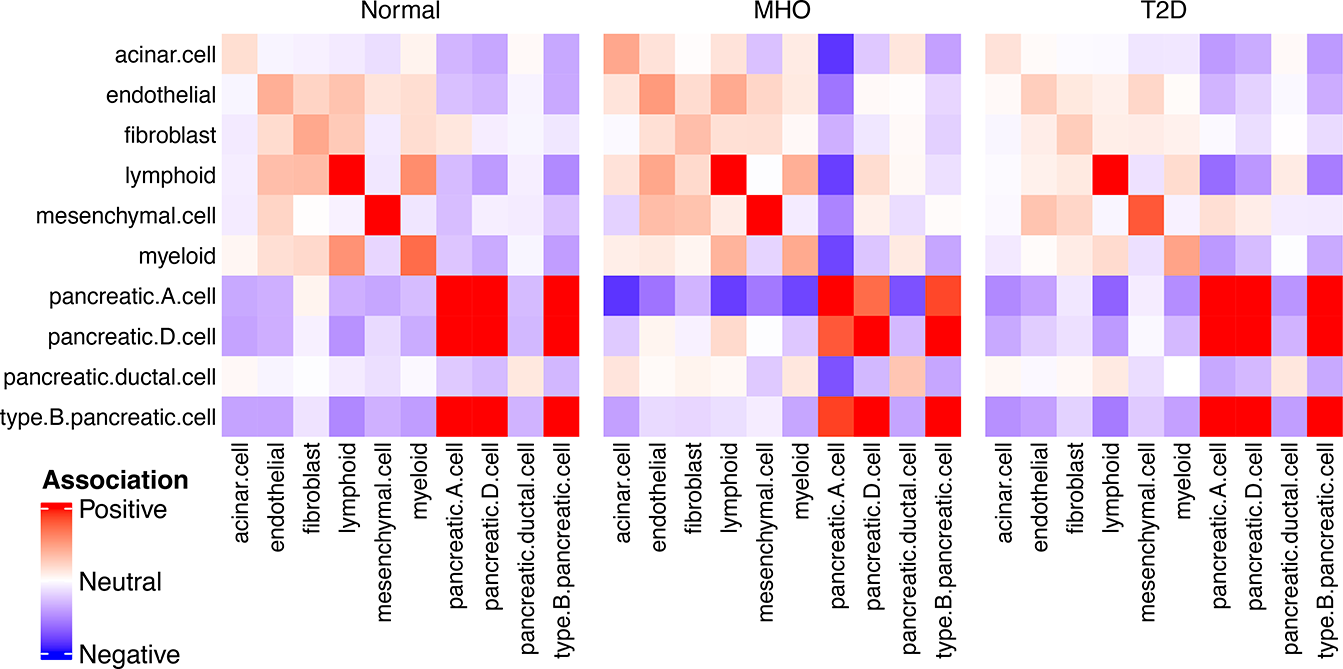
Frequency of cell neighbor association changes in the islet with the metabolic state. Association pairs between cell type (components listed on x and y axes) within islets shifts for alpha cells in MHO and T2D compared to normal individuals. In particular, alpha cells (“pancreatic.A.cell” in figure) are less associated (blue) with acinar, fibroblast, lymphoid, myeloid, and pancreatic ductal cells in MHO and T2D patients compared to normal while delta cells (“pancreatic.D.cell” in figure) were more strongly associated (red) with lymphoid cells in the MHO population but not in normal or T2D populations.

One striking finding that was similar between our data and that of Wu et al.^17^ was that alpha cells appear to be more closely associated with pancreatic acinar cells in islets from normal control patients than in islets from MHO donors (**Fig. 3**). We therefore quantified the average distances between alpha cells and acinar cells in islets from the three patient groups (**Fig. 4A**). In islets from non-obese, non-diabetic patients, alpha cells were on average 0.018 +/− 0.001 mm away from acinar cells. However, the average distance between alpha cells and acinar cells in islets collected from MHO patients was significantly higher (0.054 +/− 0.007 mm, *p* < 0.0004). In the setting of T2D, the average distance between the two cell types returned to normal baseline values (0.020 +/− 0.001 mm). Although islets contain small numbers of acinar cells within their perimeter^26^, the greater distance between alpha cells and acinar cells in MHO islets does not account for the changes in the overall numbers of acinar cells within the islets (**Fig. 4B**). Thus, our findings suggest the changes specifically in the population of alpha cells organized in the periphery of the islets. We performed differential expression (DE) analysis to explore potential transcriptomic determinants of this observation. Gene expression in alpha cells oriented close to acinar cells was compared to alpha cells that were not spatially located in close proximity to acinar cells (**Fig. 4C, E, G**) and we observed distinctive DE patterns between the three patient groups (**Supplemental Table 4**).

**Figure 4:**
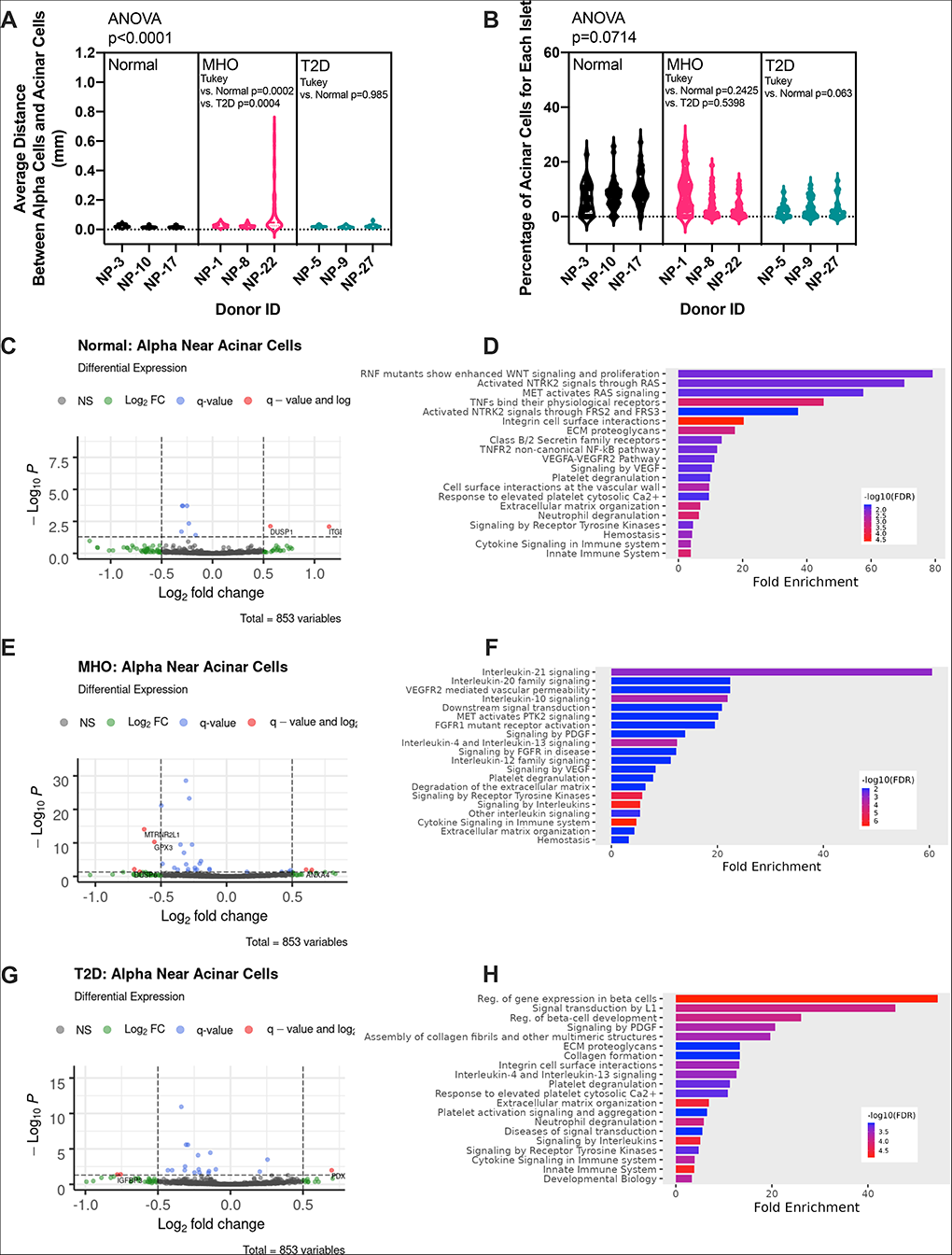
Alpha cells spatially oriented around the periphery of islets exhibit differences in gene expression across groups. A) Violin plots representing the average distances between alpha cells and acinar cells in each islet for individual donors in each experimental group. Islets from MHO donors exhibited significantly increased inter-cell distances between alpha and acinar cells compared to normal and T2D donors. No significant differences between islet acinar cell numbers were observed between groups (B). Volcano plots of differential expression between alpha cells near (right wing) versus far (left wing) from acinar cells in normal (C), MHO (E), and T2D (G) populations. A significance threshold of −log10(0.05) was used (5% of the genes called as significant are false positive). Pathway analysis within the reactome database for enriched genes in normal (D), MHO (F), and T2D (H) demonstrated a shift to interleukin signaling in the MHO population and regulators of beta cell gene expression in the T2D population.

Reactome pathway analysis (**Fig. 4D, F, H**) revealed a high degree of inflammatory cytokine signaling in peripheral alpha cells in islets from MHO patients (**Fig. 4F**). In particular, DE of genes signifying interleukin-21 (IL-21) and IL-10 family (including IL-20) signaling were enriched to a remarkable degree. IL-21 was previously shown to be essential for the development of autoimmune diabetes mellitus in NOD mice^27^; however, the role of IL-21 specifically in alpha cells has not been reported. IL-21 has been also shown to exert opposing effects depending on the presence or absence of co-stimulatory factors present in the affected microenvironment^28^. For example, in the absence of co-factors, IL-21 induces apoptosis in B lymphocytes while in the presence of IL-4, IL-21 promotes antibody production and B cell differentiation into memory cells or plasmablasts^28^. IL-20 is a member of the IL-10 family of cytokines that plays an important role in epithelial cell innate defense and repair^29^. IL-20 in particular has been shown to enhance cell proliferation^29^ and, in islets, IL-20 subfamily members restore insulin secretion and sensitivity^29^. In contrast, peripheral alpha cells in islets from patients with T2D exhibit enhancement of DE genes associated with the regulation of beta cell gene expression and beta cell development (**Fig. 4H**), suggesting conversion of peripheral alpha cells to beta cells in this context. From these data, we hypothesize that, in the setting of obesity, inflammation, lipid accumulation, and insulin resistance drive islet expansion. Given the increased diameter of islets, the alpha cells oriented around the periphery of the islets would potentially be exposed to higher levels of cytokines than alpha cells located in the core of the islets. Although continued cytokine exposure eventually contributes to reduced beta cell mass, IL-21 and IL20/IL-10 family signaling in peripheral alpha cells could promote alpha cell survival, and, perhaps, differentiation or de-differentiation^28,29^. Thus, following progression to T2D and contraction of islet area (**Fig. 2F,G**), surviving peripheral alpha cells could de-differentiate toward a beta cell phenotype in a further effort to protect against loss of insulin secretion.

In summary, CosMx Spatial Molecular Imaging of human pancreatic tissues derived from normal, MHO, and T2D patients reveals that, consistent with previous reports, obesity is associated with islet area expansion and that the percentage of beta cells within individual islets is reduced in both obesity and T2D. Our data add the novel findings that in the setting of obesity without overt T2D (MHO), lymphoid cells are preferentially oriented in close proximity with delta cells (**Fig. 3**) and that alpha cells are oriented farther away from acinar cells in islets from MHO patients. The data collected through SMI are therefore useful for confirmation of previous studies and testing of discrete hypotheses, as well as for novel hypothesis generation. Limitations of our study include small sample sizes (3 individuals per group), use of mRNA expression alone rather than the combination with protein expression to identify cell types, and previously unvalidated cell delineation protocols. In spite of these limitations, further utilization of this and related spatial technologies^16^ for analysis of molecular changes in islets and in the surrounding acinar cells are expected to provide novel insights into the mechanisms underlying the development of metabolic diseases including diabetes mellitus.

## METHODS

### Tissue Collection and Processing

Tissues from the tail of the pancreas were collected from organ donors through a collaboration with Mid-America Transplant. The Saint Louis University Internal Review Board determined that because the samples are from deceased individuals and are not associated with identifying information, the studies described herein do not constitute human subjects research. Pancreas tissues were immediately placed in formalin following removal. The post-mortem interval between physiologic death and tissue fixation was less than 30 minutes for all patients. Donor characteristics are detailed in **Supplemental Table 1**. Formalin-fixed, paraffin-embedded (FFPE) tissue sections were prepared for CosMx SMI profiling as described in He et al^11^. Briefly, five-micron tissue sections on VWR Superfrost Plus Micro slides (cat# 48311-703) were baked overnight at 60°C, then prepared for *in-situ* hybridization (ISH) by deparaffinization and heat-induced epitope retrieval (HIER) at 100°C for 15 minutes using ER1 epitope retrieval buffer (Leica Biosystems product, citrate-based, pH 6.0) in a pressure cooker. Following HIER, tissue sections were digested with 3 μg/ml Proteinase K diluted in ACD Protease Plus at 40°C for 30 minutes. Tissue sections were washed twice with diethyl pyrocarbonate (DEPC)-treated water (DEPC H_2_O) and incubated in 1:2,000 diluted fiducials (Bangs Laboratory) in 2X SSCT (2X saline sodium citrate, 0.001% Tween-20) solution for 5 min at room temperature in the dark. Excess fiducials were rinsed from the slides with 1X phosphate buffered saline (PBS) and tissue sections were fixed with 10% neutral buffered formalin (NBF) for 5 min at room temperature. Fixed samples were rinsed twice with Tris-glycine buffer (0.1M glycine, 0.1M Tris-base in DEPC H_2_O) and once with 1X PBS for 5 min each before blocking with 100 mM *N*-succinimidyl (acetylthio) acetate (NHS-acetate, ThermoFisher) in NHS-acetate buffer (0.1M NaP, 0.1% Tween PH 8 in DEPC H_2_O) for 15 min at room temperature. The sections were then rinsed with 2X saline sodium citrate (SSC) for 5 min and an Adhesive SecureSeal Hybridization Chamber (Grace Bio-Labs) was placed over the tissue. NanoString^®^ ISH probes were prepared by incubation at 95°C for 2 min and placed on ice, and the ISH probe mix (1nM 980 plex ISH probe, 10nM Attenuation probes, 1nM SMI-0005 custom, 10nM SMI-0005A1, 2nM SMI-0005A2, 1X Buffer R, 0.1 U/μL SUPERase•In™ [Thermofisher] in DEPC H_2_O) was pipetted into the hybridization chamber. The hybridization chamber was sealed to prevent evaporation, and hybridization was performed at 37°C overnight. Tissue sections were rinsed of excess probes in 2X SSCT for 1 min and washed twice in 50% formamide (VWR) in 2X SSC at 37°C for 25 min, then twice with 2X SSC for 2 min at room temperature and blocked with 100 mM NHS-acetate in the dark for 15 min. A custom-made flow cell was affixed to the slide in preparation for loading onto the CosMx SMI instrument.

### CosMx SMI-Based Target Detection

RNA target readout on the CosMx SMI instrument was performed as described in He et al.^11^ Briefly, the assembled flow cell was loaded onto the instrument and Reporter Wash Buffer was flowed to remove air bubbles. A preview scan of the entire flow cell was taken, and 18-23 fields of view (FOVs) were placed on the tissue to match regions of interest identified by H&E staining of an adjacent serial section. RNA readout began by flowing 100 μl of Reporter Pool 1 into the flow cell and incubation for 15 min. Reporter Wash Buffer (1 mL) was flowed to wash unbound reporter probes, and Imaging Buffer was added to the flow cell for imaging. Nine Z-stack images (0.8 μm step size) for each FOV were acquired, and photocleavable linkers on the fluorophores of the reporter probes were released by UV illumination and washed with Strip Wash buffer. The fluidic and imaging procedure was repeated for the 16 reporter pools, and the 16 rounds of reporter hybridization-imaging were repeated multiple times to increase RNA detection sensitivity. After RNA readout, the tissue samples were incubated with a 4-fluorophore-conjugated antibody cocktail against CD298/B2M (488 nm), PanCK (532 nm), Glucagon (594 nm), and Insulin (647 nm) proteins and DAPI stain in the CosMx SMI instrument for 2 h. After unbound antibodies and DAPI stain were washed with Reporter Wash Buffer, Imaging Buffer was added to the flow cell and nine Z-stack images for the 5 channels (4 antibodies and DAPI) were captured.

### Data Processing and Analysis

Individual cells were typed using Insitutype^30^. Using reference profiles^31^ for gene expression in immune and pancreatic tissue, cell types were assigned with a Bayes classifier based on the observed transcript molecules for each segmented cell. Dense clusters of endocrine cell types (α, β, and δ cells) were identified using the DBSCAN^32^ algorithm. A convex hull around these clusters defined the Islets of Langerhans cross-sections. Cell type co-localization evaluated Ripley’s K-function^33^ across different cell types with a range of radii from zero to 100 μm. Differences across this range between observed and theoretical value for Poisson process are summarized in heatmaps for each tissue type. A negative binomial mixed-effects model per tissue type (Normal, MHO, T2D) with patient sample as a random effect compared differential expression within cell types based on whether cells were near (less than 20 μm) or not near another cell type of interest. Reactome pathway analysis was performed using ShinyGO (http://bioinformatics.sdstate.edu/go/)^34^.

## Supporting information

Supplemental Table 1: Donor Characteristics

Supplemental Table 2: Gene Panels

Supplemental Video 1: Normal Islet

Supplemental Video 2: MHO Islet

Supplemental Video 3: T2D Islet

## AUTHOR CONTRIBUTIONS

G.R.K. designed experiments, collected and prepared tissues for SMI, designed analyses of datasets, and edited the manuscript. E.E.K. performed SMI, designed analyses of datasets, and edited the manuscript. D.R. processed the datasets, assisted in design of and performed analyses of the datasets, assembled figures, and edited the manuscript. W.K.S. assisted in experimental design, advised on analyses, and edited the manuscript. G.L.C.Y. designed experiments, designed analyses, assembled figures, performed statistical analyses, and drafted the manuscript.

## ACKNOWLEDGEMENTS

The authors would like to acknowledge Mid-America Transplant for their collaboration in obtaining pancreas tissues from the donors described in this manuscript. Without their assistance, and the generosity of the donors and their families, this study would not be possible.

## SUPPLEMENTAL INFORMATION

**Supplemental Table 1**: Patient Characteristics

**Supplemental Table 2**: Gene Target List

**Supplemental Table 3**: Quality Control and Run Metrics

**Supplemental Table 4**: Differential Expression

**Supplemental Video 1**: Resolution of Normal Islet from UMAP to Real Space

**Supplemental Video 2**: Resolution of MHO Islet from UMAP to Real Space

**Supplemental Video 3**: Resolution of T2D Islet from UMAP to Real Space

