## Supplementary figures and images for "Spatial molecular imaging of the human type 2 diabetic islet"

### Supplemental Video 1: Normal Islet

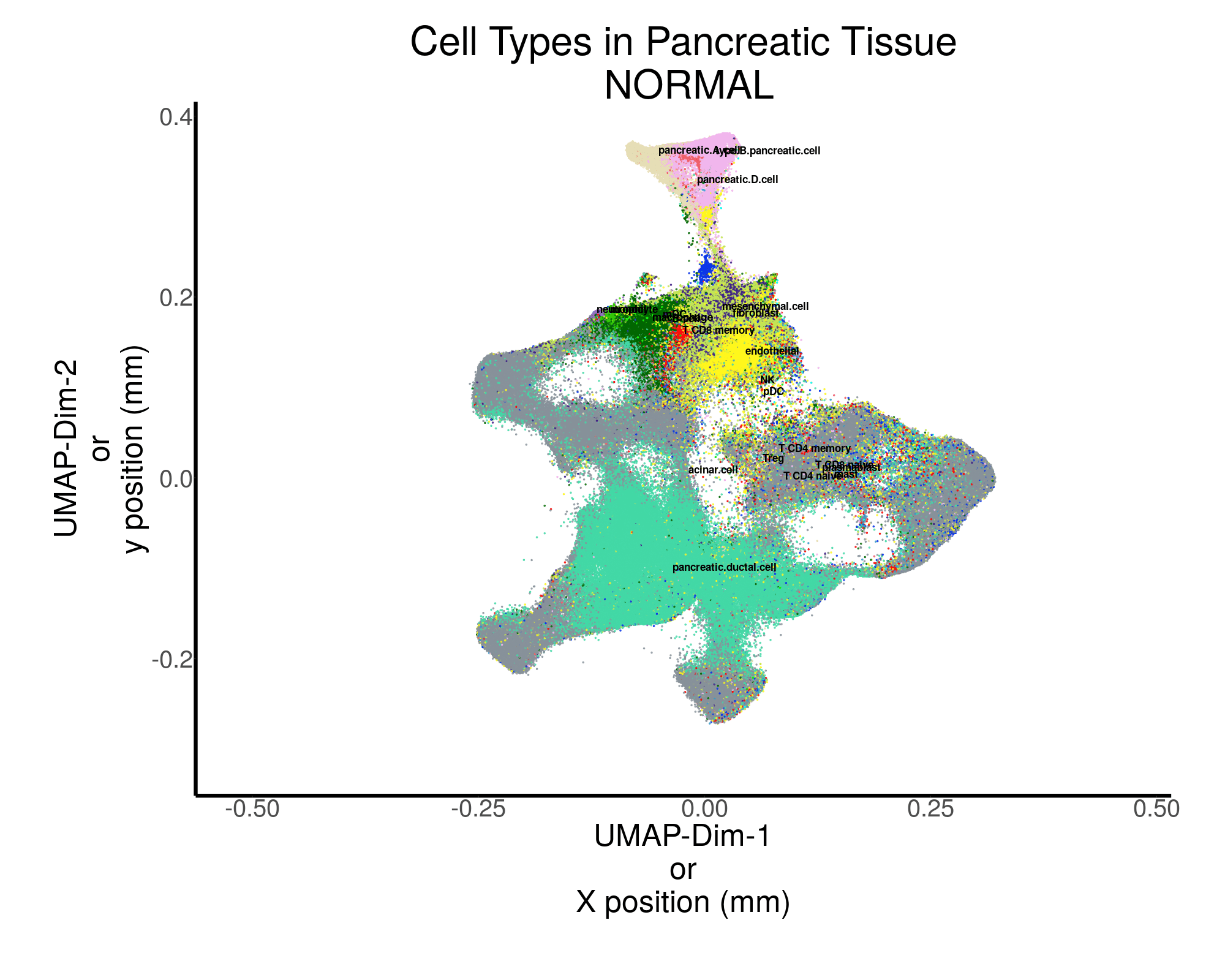

### Supplemental Video 2: MHO Islet

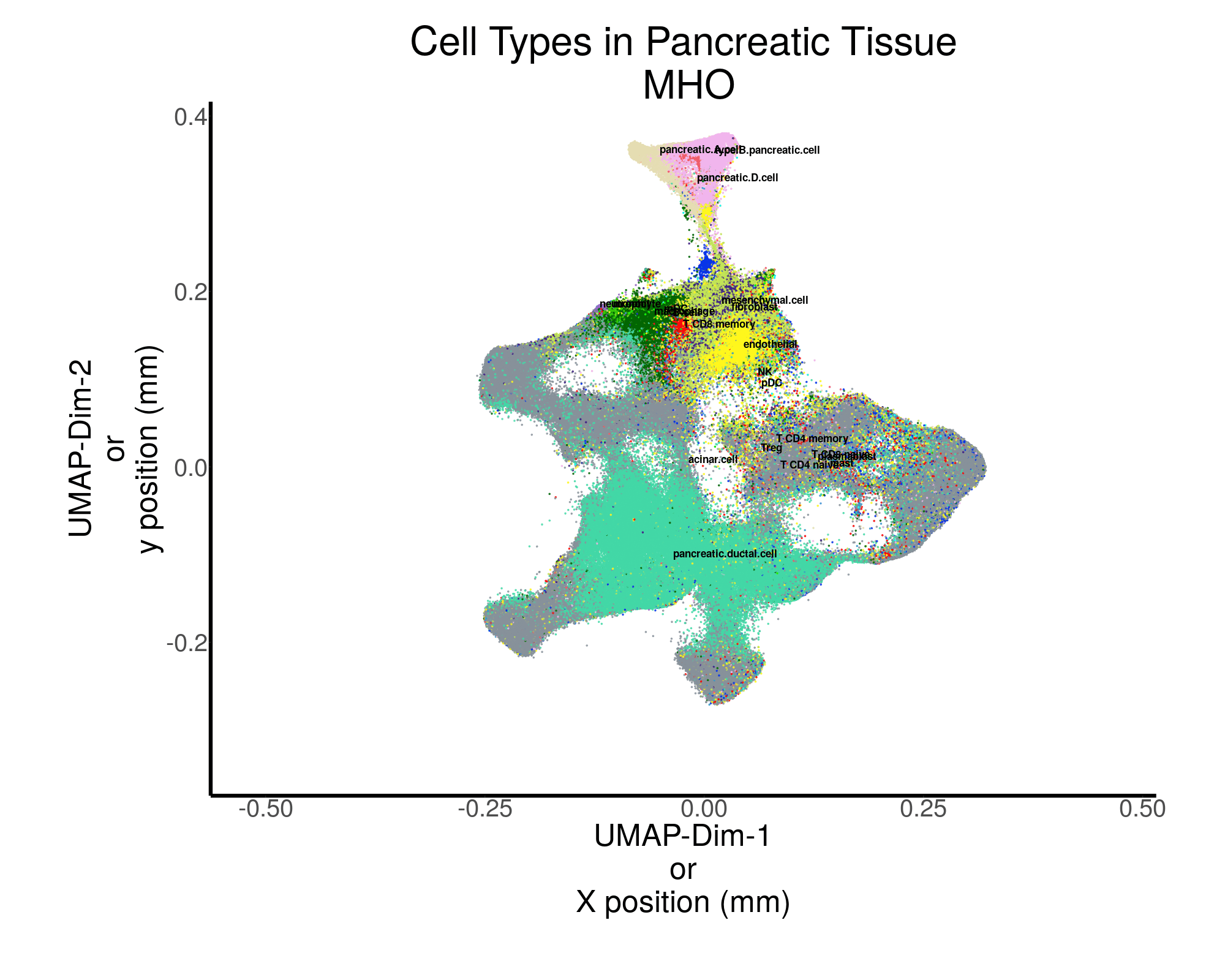

### Supplemental Video 3: T2D Islet

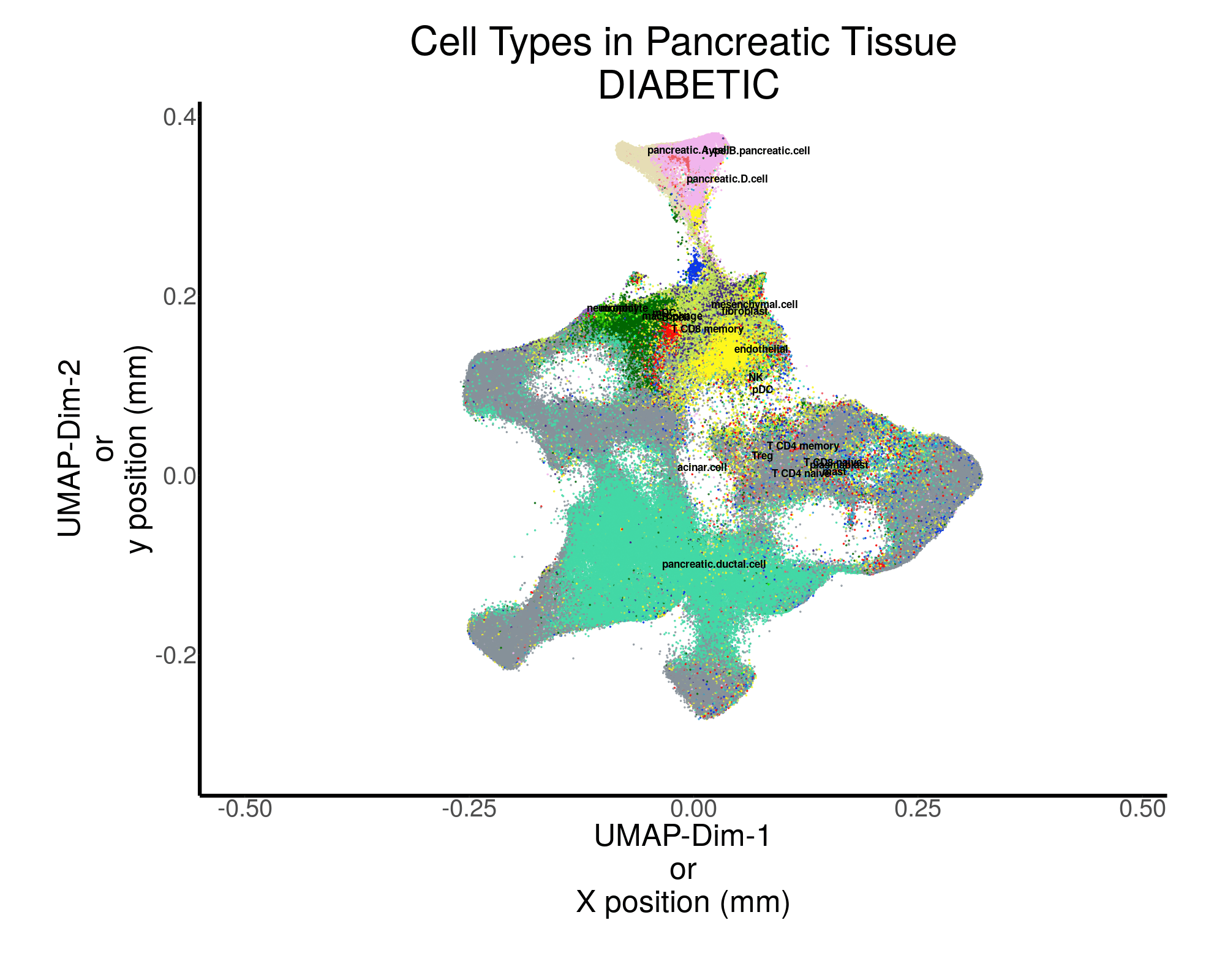
